# Outsourced hearing in an orb-weaving spider that uses its web as an auditory sensor

**DOI:** 10.1101/2021.10.17.464740

**Authors:** Jian Zhou, Junpeng Lai, Gil Menda, Jay A. Stafstrom, Carol I. Miles, Ronald R. Hoy, Ronald N. Miles

## Abstract

Hearing is a fundamental sense of many animals, including all mammals, birds, some reptiles, amphibians, fish, and arthropods. The auditory organs of these animals are extremely diverse in anatomy after hundreds of millions of years of evolution, yet all are made up of cellular tissues and are morphologically part of bodies of animals. Here we show hearing in the orb-weaving spider, *Larinioides sclopetarius* is not constrained by the organism’s body but is extended through outsourcing hearing to its extended phenotype, the proteinaceous, self-manufactured orb-web. We find the wispy, wheel-shaped orb-web acts as a hyperacute acoustic “antenna” to capture the sound-induced air particle movements that approach the maximum physical efficiency, better than the acoustic responsivity of all previously known eardrums. By sensing the motion of web threads, the spider remotely detects and localizes the source of an incoming airborne acoustic wave such as those emitted by approaching prey or predators. By outsourcing its acoustic sensors to its web, the spider is released from body size constraints and permits the araneid spider to increase its sound-sensitive surface area enormously, up to 10,000 times greater than the spider itself. The spider also enables the flexibility to functionally adjust and regularly regenerate its “external eardrum” according to its needs. The “outsourcing” and “supersizing” of auditory function in spiders provides unique features for studying extended and regenerative sensing, and designing novel acoustic flow detectors for precise fluid dynamic measurement and manipulation.

## Introduction

During the water-to-land transition, animals have gone through dramatic challenges in aerial hearing (1, 2). To effectively detect weak, distant airborne sound, terrestrial vertebrates and some invertebrates have evolved the tympanic eardrums which are very sensitive to the pressure component of sound (1, 3). Alternatively, some arthropods, especially those of miniscule size, have evolved pendulum-like, long wispy filaments to detect the velocity component of sound (4–6). While the auditory organs of different animals are extremely diverse in anatomy after hundreds of millions of years of evolution (7, 8), they are all organs of cellular origin and are morphologically part of bodies of animals.

Spiders are among the oldest land animals, with a fossil record dating back to the Devonian Period (around 380-million years ago) (9). All spiders produce silk, a biomaterial that can be stronger than steel in strength-to-weight ratio yet extremely flexible (10), owing to its exceptional material properties. When woven into a broad latticework, a web can serve as a net for capturing prey that fly or walk into it (11–13). We previously showed that a single strand of nano-dimensional spider silk can move with a velocity very close to that of the surrounding air particle movements, with the maximum physical efficiency from infrasound to ultrasound, despite the low viscosity and low density of air (14). Here we show that the highly responsive aerodynamic property of silk fibers are woven and stretched into diaphanous orb-web can function as a huge acoustic “antenna,” which allows the spider to efficiently detect faint airborne sound from a distant source. This outsourced orb-web “eardrum” as an extended phenotype beyond the body operates on a very different principle from the much smaller auditory organs of all other animals.

## Results and Discussion

We have found that the orb-weaving spiders detect and localize distant airborne sound (Fig. 1, Movie S1, Movie S2). Spiders used in this study are *Larinioides sclopetarius,* a familiar araneid related to the species celebrated by E.B. White, in “Charlotte’s Web”. All spiders were field-collected in Vestal, N.Y. and kept in laboratory conditions, where they spontaneously spin orb-webs within wooden frames (30 cm × 30 cm × 1 cm). We videotaped spider behavioral reactions (N=60 spiders) to airborne acoustic tones within a spacious closed anechoic room with a 60 fps video camera, in 2 different auditory configurations (see *SI Appendix,* Fig. S1, Table S1): 1) normal-incident sound waves to the orb-web plane, emitted from a frontally-positioned loudspeaker, 3.0 m away to the spider; and 2) oblique-incident directional sound waves, emitted from loudspeakers placed at 45 degrees to the left and right of the spider in azimuth, 0.5 m away. Before making any measurements, we ensured that the spiders were undisturbed and resting naturally within the hub region. Each spider was acoustically stimulated only once per trial. The behavior of each spider was video-monitored for 5 seconds after initiating the stimulus tone. Its behavior in the 5 seconds preceding acoustic stimulation served as the silent control period (0 dB). Within our sealed chamber there were no uncontrolled airflows or substrate vibrations (see Methods and *SI Appendix,* Fig. S2) to disturb the spider; the airborne stimuli from the speakers were the only source of web vibrations.

**Fig. 1.**
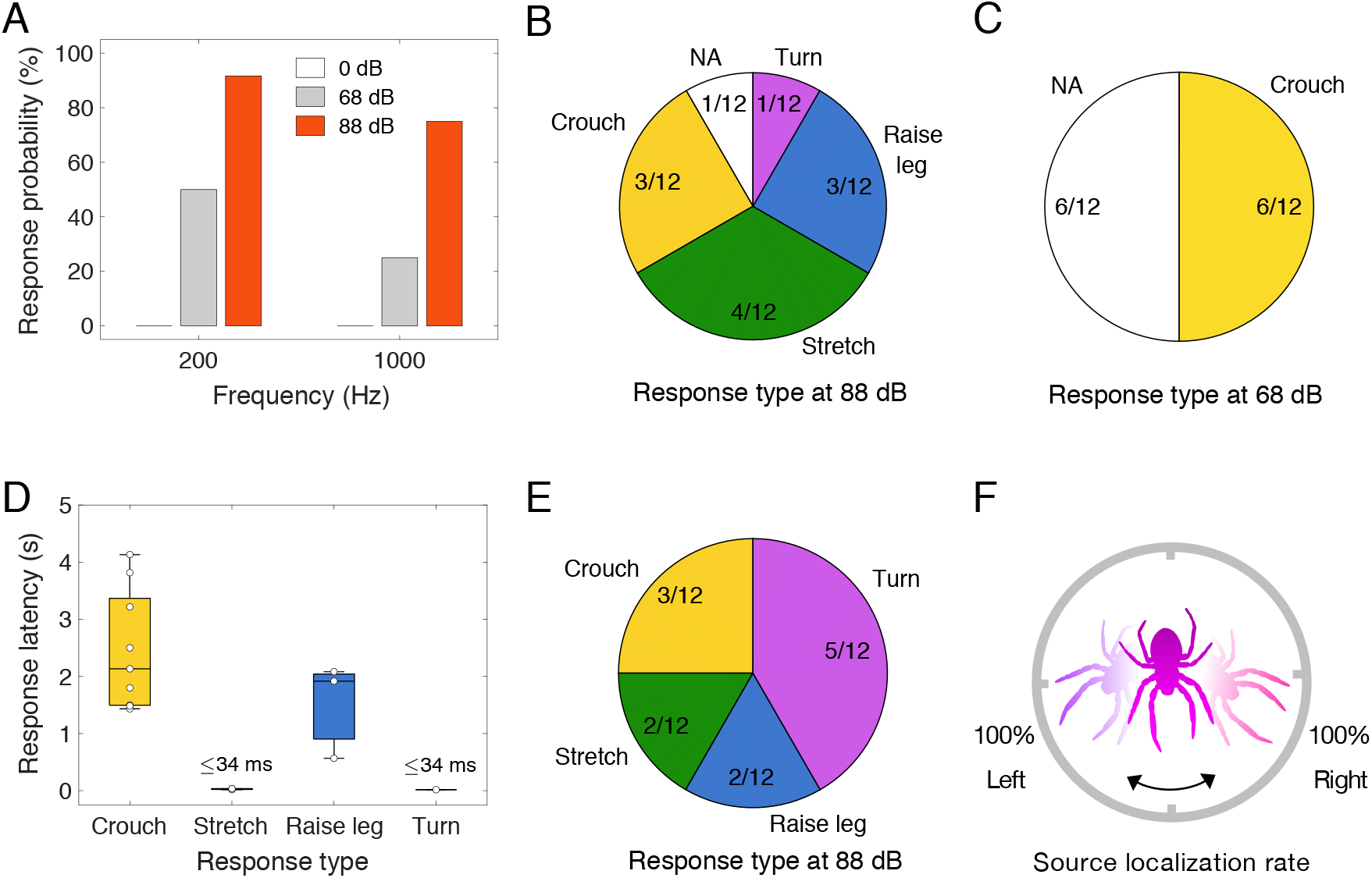
Adaptive behavioral responses made by orb-weaving spiders stimulated by remote airborne sound. Two acoustic hearing conditions of airborne tone stimulation were investigated *(SI Appendix,* Fig. S1): 1) normal-incident sound (*A*-*D*, see Movie S1), generated by a loudspeaker, 3.0 m distance to the spider, with a propagated direction perpendicular to the plane of the orb-web; 2) oblique-incident sound (*E* and *F*, see Movie S2), from one of two identical loudspeakers, 0.50 m distance, randomly placed to the left and right side of the spider, 45° in azimuth. (*A*) Percentage of spider response (4 groups, N=12 spiders in each group) to normal-incident acoustic tones of 3 s duration. (*B-D*) Behavioral response categories and response latencies (each value, median, interquartile, and range) to 200 Hz tones. NA represents no behavioral response in the pie charts. (*E*) Response (N=12 spiders) to oblique-incident acoustic tones (200 Hz, 88 dB). (*F*) Of all turning responses, percentage of turning towards the randomly assigned direction of a sound source.

The responsiveness and sensitivity of spiders to relatively distant (3.0 m) normal-incident acoustic waves are shown in Fig. 1*A-D*. Four distinct behavioral responses, in the form of rapid adaptations in body posture, were provoked by acoustic stimulation under 4 combinations of tone frequency (200 Hz and 1000 Hz) and sound pressure level (SPL, a “soft” 68 dB and an “intense” 88 dB). We assigned 48 spiders to 4 groups (N=12 spiders in each group). The duration of each tone stimulus, at a given SPL and frequency, was 3 seconds. Of 12 individuals tested in each group, 6/12 and 11/12 responded to the 200 Hz tones at 68 dB and 88 dB respectively, 3 /12 and 9/12 responded to the 1000 Hz tones to 68 dB and 88 dB sound levels, respectively, but none responded during silent controls. We identified several categories of intensity-dependent behaviors to 200 Hz tones. At high 88 dB levels (Fig. 1*B*, *SI Appendix,* Movie S1), body responses consisted of 1) *crouching* (pulling the web strands slightly with legs), 2) *stretching-out* or *flattening of the body* (rapidly extending all 8 legs outward from the cephalothorax), 3) *foreleg displaying* (lifting the forelegs into the air), and 4) *turning* (abruptly changing the direction of the body). The response to the 88 dB stimuli was complex and variable—some spiders responded sequentially with several behaviors, so we only recorded the spider’s initial response for labeling the behavior. Importantly, the only behavioral response to the low intensity stimulation, at 68 dB was *crouching* (Fig. 1*C*). Orb-weaving spiders were previously shown to be able to detect the nearby (3-4 cm) flies buzzing in the air (15), which is a near-field effect but not hearing at distances that would qualify as far-field hearing at the 3 m distances demonstrated in our present study. The sound pressure level of the biologically relevant information depends on the source distance to the spiders. According to the inverse square law, the sound pressure level drops by 20 dB for every 10 times increase in source distance, for example an 88 dB sound source at 1 m distance would be 68 dB at 10 m distance. Potential predators and prey such as birds, frogs, and crickets can produce loud sound at remote distance. The sound pressure level can be even larger than 80 dB after propagating in air for 10 m (*SI Appendix,* Table S2). Given a hearing threshold lower than 68 dB, orb-weaving spiders should be able to early detect predators and prey at a distance more than 10 m away.

We also observed that spiders can localize the direction of airborne sound accurately. Active and rapid turning movements toward the source speaker were observed when the speaker location was shifted from normal to oblique incident (45°, L/R) in azimuth (Fig. 1*E, F*, *SI Appendix,* Movie S2). Unlike normal-incident soundwaves that arrive at the orb-web plane simultaneously, oblique-incident soundwaves arrive with brief delays, creating directional acoustic cues such as time, amplitude and phase differences at different locations of the orb-web (16). Of 12 individuals tested for directional hearing, all (12/12, Fig. 1*E*) responded to oblique-incident 200 Hz tones at 88 dB, similar to the high responsivity (11/12, Fig. 1*B*) to the normal-incident sound at 88 dB from the front. However, more spiders (5/12, Fig. 1*E, F*) responded to the directional oblique-incident sound by turning towards the stimulating speaker, compared to only 1 of 12 spiders (Fig. 1*B*) that turned in response to normal-incident sound.

Having found that *L. sclopetarius* spiders exhibit behavioral responses by detecting and localizing airborne sound, we then investigated the physical properties of the web as an acoustic antenna by measuring the mechanical response of the web to direct acoustic stimulation using sounds that are salient to spider hearing (Fig. 2, Movie S3). Using Doppler vibrometry, we measured sound-induced out-of-plane web motion for the web alone and with live spiders resting in the web’s hub. The web was separated from a loudspeaker 3.0 m away and aligned so that the direction of sound propagation was perpendicular to the plane of the orb-web, creating a normal-incident planar sound wave *(SI Appendix,* Fig. S1*E*, *F*). We broadcast sinusoidal test tones ranging from 100 Hz to 10,000 Hz. The air particle velocity component, u(t), of a sound wave was computed from the measurement of the pressure p(t) using a calibrated pressure-sensitive microphone near, but not touching the web, where u(t)=p(t)/ρ_0_c, ρ_0_ is the density of air, c is the speed of sound in air (17). Fig. 2*A* shows an example of the measured orb-web response to 200 Hz acoustic signals, in which the orb-web follows the air particle motion with almost full fidelity and maximum physical efficiency (i.e. V_web_/V_air_~1), better than the acoustic responsivity of all known eardrums (14, 18). Similarly, at other measurement frequencies, orb-webs responded effectively to airborne acoustic signals, a bandwidth encompassing the sounds produced by potential prey and predators, such as insects and birds (Fig. 2*C, E*, *SI Appendix,* Fig. S3 and S4, and Audio S1). The differences in velocity between the spider body and the web (Fig. 2*D, E*) suggest that mechanical strain is actively induced on the spider’s legs when stimulated by airborne sound. In spiders, vibrational signals such as faint acoustic stimuli are presumably detected by the strain-sensitive lyriform organs located in spider legs (19–21).

**Fig. 2.**
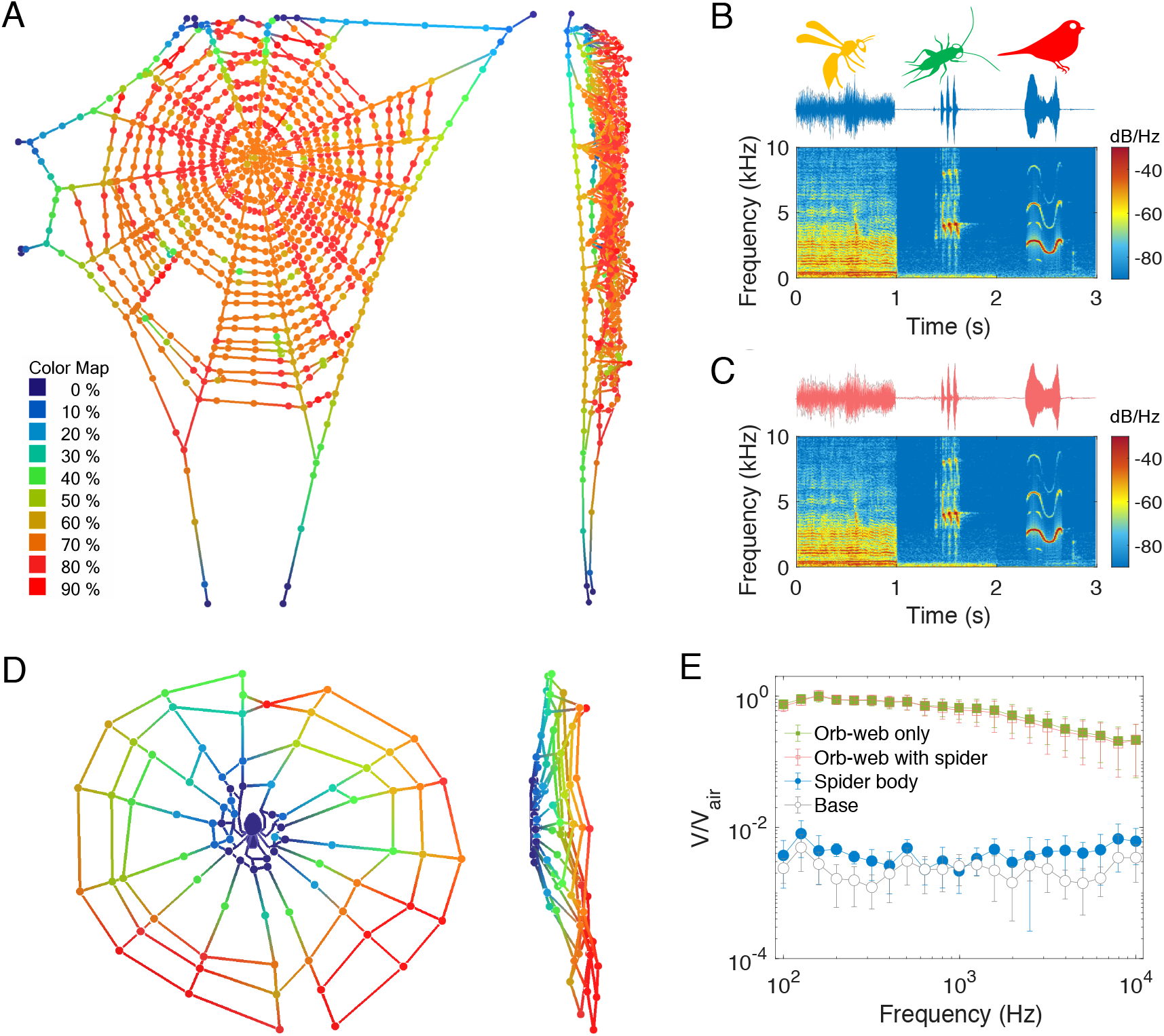
Spider orb-web is an enormous, reconfigurable, regenerative, and highly sensitivity acoustic antenna. (*A-E*) Orb-web responses to remote normal-incident sound, generated by a loudspeaker, 3.0 m distance to the spider. (*A*) Out-of-plane motion of a complete orb-web induced by a 200 Hz steadystate sound field (see Movie S3). The colored heat map represents the amplitude ratio of the web thread velocity V to the air particle velocity V_air_, e.g. 90% represents V/V_air_=0.9~1.0. (*B* and *C*) Time-domain traces and spectrograms of the airborne acoustic signal (34) measured by a pressure microphone (*B*), and the signal-induced web motion measured by a laser vibrometer (*C*) at the hub web position as shown in (*A*). The audio clip of the measured web motion is provided as Audio S1. The acoustic signal spans a wide range of frequencies (100 Hz-10000 Hz) including hymenopteran wing-beats, cricket chirpings, and bird songs. (*D*) Heat map depiction of out-of-plane motion of web containing a live spider, induced by a 200 Hz sound field (Movie S3). Color coding of (*D*) same as (*A*). (*E*) Statistic frequency responses of the spider orb-webs (N=12) to airborne sound. Individual measurement (1 of 12) contains 1 location on a spider body, 1 location on an orb-web frame base, and 4 locations (up, down, left, and right) on radial threads of an orb-web in radial distance of 5 cm away from its hub, without and with the spiders resting in the hub webs. Error bars show one standard deviation (SD).

It is important to note that some spider species and insects can detect air particle velocity with pendulum-like, long, wispy cuticular hair receptors (4–6, 18). It is also widely known that spiders can sense movements of the web when a vibrating stimulus is applied directly (touching) to the web silk (12, 13, 22). To determine whether the orb-weaving spider *L. sclopetarius* is detecting highly circumscribed sound-induced web movements or is directly detecting some fluctuating acoustic quantity of the air (such as pressure or velocity) through some unidentified mechanosensor, we stimulated only small, focal regions of the web while ensuring that any airborne acoustic signal would have such a low amplitude that it would not be detectable by the spider, perched in the center of the web, distant from such a focal acoustic stimulus.

To accomplish this, we used a miniature speaker (dimensions 15 mm × 11 mm × 3 mm) as a focal sound source (Fig. 3*A*). The speaker was positioned 50 mm in radial distance away from the spider, that was resting in the web’s hub. By aligning the small speaker as near as possible to the web without actually touching it, we created a localized “near-field” sound causing oscillations in air particle velocity, from the mini-speaker, but which rapidly decayed with distance as it propagates through the air (Fig. 3*B, C*). We showed that the near-field airborne stimulation generated by the mini-speaker attenuated quickly with distance, and fell well below the spider’s detection threshold after it spread to distantly perched spider (<50 dB, see Fig. 3*B, C* and *SI Appendix,* Fig. S5). However, the out-of-plane web movements induced by the airborne sound created by the mini-speaker attenuated less, so that vibrational signals were transmitted to the spider. Results show that the spiders perceived minute localized web vibrations at extremely low intensity levels (Fig. 3*D, E*). Of 12 individuals, 4 responded to 200 Hz web vibration tones of 3 s duration with equivalent SPL ≤ 68 dB (V_rms_ ≤ 0.12 mm/s). Spiders responded to these minute web vibrations by *crouching,* just as they respond to airborne stimuli at 68 dB from a loudspeaker 3 m away (Fig. 1*C, D*). The behavioral response of spiders to minute web vibrations induced by the focal airborne sound confirms their abilities to perceive airborne acoustic signals solely by detecting web movements. In earlier work, Uetz et al. hypothesized and tested that nearby (up to 20 cm) acoustic stimulation might be an “early warning” channel against predators (23). The web-enabled hearing with threshold lower than 68 dB should allow the spider to early detect and localize predators and prey at a distance more than 10 m away (*SI Appendix,* Table S2).

**Fig. 3.**
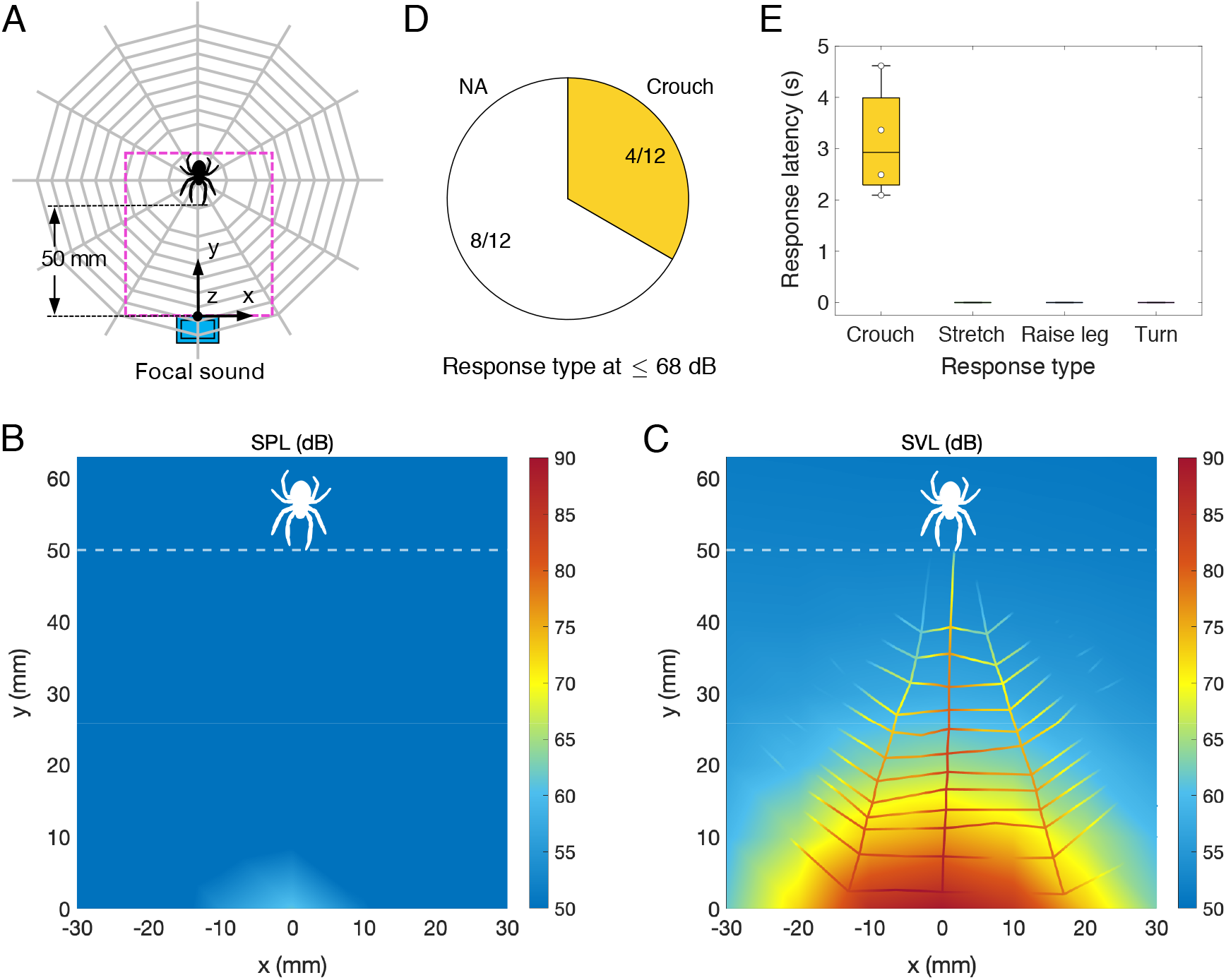
Spider behavioral responses to orb-web movements stimulated by focal airborne sound. (*A*) Schematic diagram of setup. Highly localized near-field sound was generated by a miniature speaker, placed as close to the orb-web without touching it, 50 mm distance in radial distance from a spider resting in its hub web. (*B* and *C*) Focal airborne sound field at 200 Hz, measured in the orb-web plane as marked by the magenta rectangular region in (*A*). The near-field sound is velocity dominant, where the sound pressure level (SPL in *B*) is almost ignorable in compared with the sound velocity level (SVL in *C*). Minispeaker airborne acoustic signals attenuate quickly and fell well below the detection threshold of spiders (see Methods) after propagating to where the spider sat (< 50 dB), while the induced out-of-plane web movements (*C*) attenuate less (≤ 68 dB). (*D* and *E)* Behavioral response categories and response latencies (each value, median, interquartile and range) to focal tones (200 Hz, equivalent SPL ≤ 68 dB, 3 s duration).

Outsourcing the acoustic sensors to its web provides the spider with the flexibility to adjust its hearing adaptively according to its needs. When an orb-web is torn or badly damaged that disrupts its radial symmetry, an orb-weaver can recover its hearing through the orb-web antenna by weaving a new one within an hour. By adjusting web geometries and pre-tensions during the web weaving, the level and tuning of the mechanical responsivity of the web threads can be both adjusted (See *SI Appendix,* Supplementary Information Text, and Fig. S6). For example, a variably-tensioned orb-web could efficiently filter out the bio-irrelevant low frequency noises which are unavoidable in the natural environment, such as the wind perturbation which has tremendous velocity and pressure amplitude than that of biorelevant acoustic signals. In principle, the multi-pedal spider can adaptively tune its hearing in real-time by behaviorally manipulating its extended “virtual eardrum”. Each of its 8 legs is endowed with sensitive vibration receptors that can be extended in all directions from the center of the web, representing “well-connected” nodes in the wheel-shaped network, consisting of smaller local nodes to interface with the web dynamics. On the one hand, the 8 legs are points of sampling for sensing, and on the other hand they have the potential to serve as feedforward controllers by adjusting postures and positions that may change directionality and sensitivity actively (24, 25). The size and shape of an orb web can be varied to meet the needs of a spider’s sensory and feeding ecology and demonstrates a remarkable level of flexibility in this surprising bioacoustic control system.

Biologists and material scientists are still discovering new properties of spider silk that can be repurposed as a biomaterial and deployed for practical human applications. Here, we demonstrate how a spider web made of nanoscale protein fibers serves as a megascale acoustic airflow sensor, contrasted sharply with all auditory organs made up cellular tissue, and necessarily subjected to body limitations. Taking advantage of the extended phenotype, the sensory surface area is up-scaled extensively, up to 10,000 times greater than the spider itself (26), much as a radio-telescope senses electromagnetic signals from cosmic sources. The acoustic function of the orb-web is analogous to an eardrum in other animals, but it senses the velocity of air particles, not its collective pressure. Spider webs are marvels of bio-architecture that greatly extend the spider’s capacity to sense and capture prey much larger than the spider itself (27, 28). The spiders also have the flexibility to tune and regenerate their hearing by manipulating the orb-webs. The new sensory modality of hearing could provide unique features for studying extended and regenerative sensing, where the orb-web functions as an integral part of the cognitive systems of a spider (28–30). The novel hearing mechanism could also presage a new generation of acoustic fluid-flow detectors in the domain of nanoscale biosensors for applications requiring precise fluid dynamic measurement and manipulation (31–33).

## Methods

### Spiders and their orb-webs

Orb-weaving spiders, *Larinioides sclopetarius,* were collected from natural habitats in Vestal, N.Y., and kept in laboratory conditions (approx. 22°C temperature, a 12 h: 12 h light-dark cycle), where they spontaneously spin orb-webs within wooden frames (30 cm × 30 cm × 1 cm) in transparent enclosures with fruit flies *(Drosophila).* After weaving the orb-webs, spiders positioned themselves to settle within the hub region of the orb-webs, as in nature. A red dim light was always turned on to enable basic visualization for setting up experiments during all light-dark cycles. Female spiders were used in all experiments. The ranges of body length and weight of spiders are 5-8 mm and 0.7-1.9 mN, respectively.

### Experimental setups

Three kinds of speaker configurations were used to create different airborne sound waves around the orb-web: 1) remote normal-incident sound wave propagating in the direction perpendicular to the plane of the web was generated by a subwoofer (Coustic HT612) and a tweeter (ESS Heil AMT1) with the crossover set at 2 kHz, placed 3.0 m away to the plane of the orb-web as shown in *SI Appendix,* Fig. S1*A*; 2) oblique-incident sound was created by two identical loudspeakers (NSM Model 5), placed 0.50 m away, 45° in azimuth on the left and right side of the spider, as shown in *SI Appendix,* Fig. S1*B*; 3) focal sound was generated by a miniature speaker (CUI CMS-15118D-L100, dimensions 15 mm × 11 mm × 3 mm), placed close to the web without touching (about 2 mm distance to the orb-web plane), 50 mm in radial distance from the spider resting in its hub web as shown in Fig. 3*A*.

The sound pressure p(t) around the orb-web was measured by a calibrated microphone (B&K 4138) placed close to the orb-web without touching. The out-of-plane motion of the orb-web was measured by a laser Doppler vibrometer (Polytec OFV-534). To measure different locations, the laser Doppler vibrometer was mounted on a precise 2-dimensional linear stage (Newport ILS250PP), controlled by a motion controller (Newport model ESP 301). The built-in camera within the vibrometer enabled the visualization of the measurement position on the orb-web. Base vibration was measured with a tri-axial accelerometer (PCB 356A11). A data acquisition system (NI PXI-1033) was used to acquire data. Spider behavioral responses were recorded using a video camera of 60 fps.

### Testing of spider behavioral response to airborne sound

The spiders’ behavioral reactions to airborne, tonal stimuli were videotaped under 3 acoustic conditions: 1) normal-incident sound waves (N=48 spiders), 2) oblique-incident directional sound waves (N=12 spiders), and 3) small focused beams of sounds (N=12 spiders). We used a single presentation of sound stimulus with duration of 3 s for the normal-incident and focused sound experiments. For the oblique-incident directional sound experiment, we used 4 successive presentations of very short sound stimulus. Individual spider was randomly subjected to one of the patterns of directional stimuli, either L+R+L+R or R+L+R+L, where L or R represents the stimulus from left or right direction with 0.3 s duration, the symbols (+) represent silent gaps between the adjacent stimulus, which are 0.2 s, 1 s, and 0.2 s respectively. Spider behavioral reactions to the successive oblique-incident sound stimuli are provided in Table S3. Detailed experimental configurations are listed in Table S1. In all three kinds of behavioral experiments, each spider was acoustically stimulated only once per trial.” Before making any measurements, we ensured that spiders were undisturbed and resting in the hub of their orb-webs. To guarantee this, after gently placing it in the testing area, the spider was left alone in the anechoic chamber for 0.5 h before playing tonal stimulation. At 88 dB stimulation, we noted that the spider’s body posture sometimes resulted in a very slight rotation, so we labeled such behavior as a *turn* only when a spider’s turn angle was unambiguously greater than 10 degrees. The response latency of an individual spider was counted from the recorded video frame number starting from the beginning of the stimuli. For all initial *turning* reactions to the oblique-incident sound, percentage of turning towards the randomly assigned direction of a sound source was given in Fig. 1F.

### Characterization of the normal-incident sound field

By placing speakers far away (3.0 m) from the spider web in our anechoic chamber (*SI Appendix,* Fig. S1*A*), we approximately created a normal-incident planar acoustic field. The direction of the propagation of the sound waves was roughly perpendicular to the plane of the orb-web. *SI Appendix,* Fig. S1*E* shows an example of the measured sound pressure level (SPL) at 200 Hz around the spider orb-web in an area of 240 mm × 240 mm with scanned gap distance 10 mm, which is considerably uniform with a SPL variation within 1 dB at different locations. The broadband SPL at all measured locations is shown in *SI Appendix*, Fig. S1*F*, which has a variation within 2 dB under the measured frequency. The sound field around the spider web can be regarded as a plane wave approximately, considering the little variation of SPL. For a plane sound wave, the air particle velocity u(t) nearby the orb-web can be determined by measuring the sound pressure p(t) according to u(t)=p(t)/ρ_0_c, where ρ_0_ is the density of air, c is the speed of sound in air. The sound pressure level (SPL) was calculated by SPL=20log_10_(P_RMS_/P_ref_), where P_RMS_ is the root mean square of the measured sound pressure p(t), P_ref_ =20 μPa is the reference pressure.

### Measurement of the airborne acoustic responsivity of the orb-webs

We characterized the airborne acoustic responsivity of the orb-webs under well-controlled normal-incident plane-wave sound field (*SI Appendix,* Fig. S1*A*). To precisely characterize the frequency responses, we used a short period of pure tones at various frequencies from 100 Hz to 10000 Hz. The measured data was then processed at each frequency with Least Squares Data Fitting to estimate the amplitude and phase of the measured airborne acoustic waves from microphone and the motion of the web threads measured by the laser vibrometer. To measure the velocity responses of a spider and its peripheral orb-web, it is crucial to keep the spider stable during the measurement process. To avoid the motion interruption from the behavioral responses, we stimulated a spider with acoustic tones (200 Hz, 88 dB, 3 s duration) before frequency measurement. After several cycles of acoustic stimulus, a spider rarely responded to the stimuli and kept stable so the measurement could be completed.

### Mapping the orb-web motion induced by airborne sound

We mapped the out-of-plane motion of the orb-web at different locations with and without the spider resting in hub web respectively, and then recreated the motion based on the measured velocity response of web threads at different locations under a certain measurement frequency f. Since the air particle velocity can be approximately regarded to be uniform in the normal-incident plane-wave sound field, the air particle velocity around all measured web threads can be expressed as u(t)=Ue^iφ^e^iωt^, where ω=2πf, U and φ are the velocity amplitude and phase of air particle motion. The time response of a measured web point in response to a steady-state airborne sound can be expressed as v_n_(t) = V_n_e^iω_n_^ e^iωt^, where V_n_ and φ_n_ are the velocity amplitude and phase of the measured web point motion. As the measured frequency responses of web threads at different locations contain the velocity amplitude as well as phase, these results can be used to create the steady-state motion of the measured objects. Figures and Movies (Fig. 1*A*, Fig. 1*D*, and Movie S3) demonstrating the out-of-plane motion of the spider and its orb-web were created by STAR 7, a commercial modal analysis software.

### Characterization of the focal sound field

We characterized the airborne signals as well as the out-of-plane transverse motion of web strands induced by the miniature speaker. Both sound pressure level (SPL, Fig. 3*B*) and sound velocity level (SVL, Fig. 3*C*) of the sound field were characterized. We scanned the acoustic pressure field by a probe microphone. The acoustic velocity field was characterized by an easily made velocity probe, constituted by the laser Doppler vibrometer and a strand of spider silk (14). The spider silk has a sub-micron diameter, 5 mm length, and was supported at its two ends loosely. By focusing the laser beam perpendicular to the longitudinal direction of the spider silk at its middle position, we measured the silk motion induced by the motion of the air particles. Before scanning of the velocity field in the orb-web plane, we confirmed that the silk velocity probe representing the air particle motion closely (i.e. V_silk_/V_air_~1) in the measured frequency, as shown by the insert figure in *SI Appendix,* Fig. S5*A*. Since velocity is a vector, we scanned the acoustic particle velocity in 3 dimensions, including V_x_, V_y_ and V_z_. The overall amplitude of velocity V (Fig. 3*C*) at a position was evaluated by 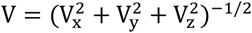. To compare between the air particle velocity and sound pressure, the sound velocity level (SVL) was calculated by SVL=20log_10_(V_RMS_/V_ref_), where V_RMS_ is the root mean square of the measured particle velocity V, V_ref_ = P_ref_/ρ_0_c is the reference velocity.

Before propagating to the location of spider, the nearfield airborne signals fell well below the detection threshold of the spiders (4, 20). The SPL (Fig. 3*B*) was below 30 dB, while the SVL (Fig. 3*C*) was lower than 50 dB (V_rms_<0.016 mm/s) after reaching to the spider. The ultralow airborne signal is even lower than the detection threshold of the jumping spider (4), which enables the best-known spider sensitivity (~65 dB) so far. Meanwhile, we never observed any behavioral response of spiders to 50 dB normal-incident airborne stimuli.

The out-of-plane transverse motion of web strands induced by the mini-speaker attenuated slower than the near-field airborne signals, so as to transmit the local vibrational signals to the spider (Fig. 3*C* and *SI Appendix,* Fig. S5*D*). Since the initial SVL generated by the mini-speaker 50 mm away from the spider is about 88 dB, and it attenuates about 20 dB after 40 mm, the equivalent SPL of the vibrational signals transmitted to the spider is less than 68 dB.

## Author Contributions

Conceptualization: J.Z, R.R.H, R.N.M; Methodology: J.Z, J.L, G.M, J.A.S, C.I.M, R.R.H, R.N.M; Investigation: J.Z, J.L, G.M, J.A.S; Formal Analysis: J.Z, J.L, G.M, J.A.S, C.I.M, R.R.H, R.N.M; Resources: C.I.M, R.R.H, R.N.M; Funding acquisition: R.N.M; Writing – original draft: J.Z, R.R.H; Writing – review & editing: J.Z, J.L, G.M, J.A.S, C.I.M, R.R.H, R.N.M; Supervision: R.R.H, R.N.M.

## Acknowledgments

We thank Charles Walcott for his expert consultation and advice. This work was supported in part by the National Institute on Deafness and Other Communication Disorders of the National Institutes of Health under Award Number R01DC017720 (to R.N.M).

## References

1. W. A. van Bergeijk, Evolution of the sense of hearing in vertebrates. Am. Zool. 6, 371–377 (1966).

2. C. B. Christensen, J. Christensen-Dalsgaard, P. T. Madsen, Hearing of the African lungfish *(Protopterus annectens)* suggests underwater pressure detection and rudimentary aerial hearing in early tetrapods. J. Exp. Biol. 218, 381–387 (2015).

3. D. Robert, J. Amoroso, R. R. Hoy, The evolutionary convergence of hearing in a parasitoid fly and its cricket host. Science 258, 1135–1137 (1992).

4. P. S. Shamble, et al., Airborne acoustic perception by a jumping spider. Curr. Biol. 26, 2913–2920 (2016).

5. G. Menda, et al., The long and short of hearing in the mosquito *Aedes aegypti*. Curr. Biol. 29, 709–714.e4 (2019).

6. C. J. Taylor, J. E. Yack, Hearing in caterpillars of the monarch butterfly *(Danaus plexippus)*. J. Exp. Biol. 222(2019).

7. R. R. Fay, L. A. Wilber, Hearing in vertebrates: a psychophysics databook, (Hill-Fay Associates, 1989).

8. J. T. Albert, A. S. Kozlov, Comparative aspects of hearing in vertebrates and insects with antennal ears. Curr. Biol. 26, R1050–R1061 (2016).

9. W. A. Shear, J. M. Palmer, J. A. Coddington, P. M. Bonamo, A devonian spinneret: early evidence of spiders and silk use. Science 246, 479–481 (1989).

10. F. G. Omenetto, D. L. Kaplan, New opportunities for an ancient material. Science 329, 528–531 (2010).

11. L. H. Lin, D. T. Edmonds, F. Vollrath, Structural engineering of an orb-spider’s web. Nature 373, 146–148 (1995).

12. W. M. Masters, H. Markl, Vibration signal transmission in spider orb webs. Science 213, 363–365 (1981).

13. C. V. Boys, The influence of a tuning-fork on the garden spider. Nature 23, 149–150 (1880).

14. J. Zhou, R. N. Miles, Sensing fluctuating airflow with spider silk. Proc. Natl. Acad. Sci. 114, 12120–12125 (2017).

15. D. Klärner, and F. G. Barth, Vibratory signals and prey capture in orb-weaving spiders *(Zygiella x-notata, Nephila clavipes;* Araneidae). J. Comp. Physiol. 148: 445–455 (1982).

16. M. A. Landolfa, F. G. Barth, Vibrations in the orb web of the spider *Nephila clavipes:* cues for discrimination and orientation. J. Comp. Physiol. A 179, 493–508 (1996).

17. R. N. Miles, “One Dimensional Sound Fields” in Physical Approach to Engineering Acoustics, R. N. Miles, Ed. (Springer International Publishing, 2020), pp. 35–52.

18. P. Fratzl, F. G. Barth, Biomaterial systems for mechanosensing and actuation. Nature 462, 442–448 (2009).

19. J. A. Stafstrom, G. Menda, E. I. Nitzany, E. A. Hebets, R. R. Hoy, Ogre-faced, net-casting spiders use auditory cues to detect airborne prey. Curr. Biol. 30, 5033–5039.e3 (2020).

20. C. Walcott, W. G. van der Kloot, The physiology of the spider vibration receptor. J. Exp. Zool. 141, 191–244 (1959).

21. F. G. Barth, Geethabali, Spider vibration receptors: Threshold curves of individual slits in the metatarsal lyriform organ. J. Comp. Physiol. 148, 175–185 (1982).

22. B. Mortimer, A. Soler, C. R. Siviour, R. Zaera, F. Vollrath, Tuning the instrument: sonic properties in the spider’s web. J. R. Soc. Interface 13, 20160341 (2016).

23. G. W. Uetz, J.A.Y. Boyle, C. S. Hieber, and R. S. Wilcox, Antipredator benefits of group living in colonial web-building spiders: the ‘early warning’ effect. Anim. Behav. 63: 445–452 (2002).

24. T. Watanabe, Web tuning of an orb-web spider, *Octonoba sybotides*, regulates prey-catching behaviour. Proc. R. Soc. Lond. B 267, 565–569 (2000).

25. K. Nakata, Attention focusing in a sit-and-wait forager: a spider controls its prey-detection ability in different web sectors by adjusting thread tension. Proc. R. Soc. B 277, 29–33 (2010).

26. M. Gregorič, I. Agnarsson, T. A. Blackledge, M. Kuntner, Darwin’s bark spider: giant prey in giant orb webs *(Caerostris darwini*, Araneae: Araneidae)? J. Arachnol. 39, 287–295 (2011).

27. M. Nyffeler, R. Altig, Spiders as frog-eaters: a global perspective. J. Arachnol. 48, 26–42 (2020).

28. H. F. Japyassú, K. N. Laland, Extended spider cognition. Anim. Cogn. 20, 375–395 (2017).

29. L. E. Miller, et al., Sensing with tools extends somatosensory processing beyond the body. Nature 561, 239–242 (2018).

30. L. R. Hochberg, et al., Neuronal ensemble control of prosthetic devices by a human with tetraplegia. Nature 442, 164–171 (2006).

31. T. Nakata, et al., Aerodynamic imaging by mosquitoes inspires a surface detector for autonomous flying vehicles. Science 368, 634–637 (2020).

32. E. K. Sackmann, A. L. Fulton, D. J. Beebe, The present and future role of microfluidics in biomedical research. Nature 507, 181–189 (2014).

33. M. Raissi, A. Yazdani, G. E. Karniadakis, Hidden fluid mechanics: Learning velocity and pressure fields from flow visualizations. Science 367, 1026–1030 (2020).

34. Sound clips: Catalog no. ML 125254, ML 208175 and ML 516988, Macaulay Library at the Cornell Laboratory of Ornithology, Ithaca, NY.

