## Supplementary information text for "Outsourced hearing in an orb-weaving spider that uses its web as an auditory sensor"

**This PDF file includes:**

Supplementary text
Figures S1 to S6
Tables S1 to S3
Legends for Movies S1 to S3
Legends for Audio S1
SI References

**Other supplementary materials include the following:**

Movies S1 to S3
Audio S1

**Supplementary Information Text**

**Comparison of acoustic flow sensing with acoustic pressure sensing.** Some spiders and insects can detect air particle velocity with pendulum-like hairs which are driven to move by the viscous force of the surrounding medium in the presence of a sound field (1, 2). The essential mechanism of this viscous force is entirely related to the physical properties of the fluid, that is, its viscosity. By eliminating the constraints of morphogenesis, orb-weaving spiders have scaled up this elemental property into a wheel-shaped network of silks into a huge web that captures sound over a large surface area (3), by up to a square meter scale (1 $m^2$), or 10,000 times greater than the spider body surface area in 1 $cm^2$ scale.

The enormously extended sound-sensitive surface area is far beyond the reach by sensing sound pressure with solid eardrums. In a scale perspective, the body size of animals limits the size of the true eardrums. In a mechanics perspective, it's not feasible to scale a thin solid eardrum up to meter-scale in the atmosphere environment due to its larger impedance, where a little bit of pressure variation (for example pressure variation induced by wind or sound waves) would destroy the eardrum. To see this, for a fixed circular membrane under uniform pressure P, the maximum stress occurs within the membrane is $\sigma=0.75P(R/t)^2$, where R and t are the radius and thickness of the circular membrane respectively (4). In natural environment, an airflow such as wind propagating perpendicular to a flat surface would induce a pressure $P=0.5C_d\rho v^2$, where v is the flow speed, ρ is the volume density of air (ρ=1.2 kg/$m^3$ at 20 °C), $C_d$ is the drag coefficient, $C_d$=1.17 for a flat circular membrane (5). For a circular membrane (A=1 $m^2$, t=1 μm) subjected only under light air breeze v=1 m/s, the wind-induced pressure can be estimated to be P=0.7 Pa (SPL=91 dB, a loudness which can also be produced by many animals such as birds and crickets), the maximum stress of the membrane is σ=167 Gpa, a value more than 100 times higher than the ultimate strength of common materials. For comparison, the tensile strength of spider silk is about 1.1 GPa (6), the high-tensile engineering steel is 1.3 GPa.

Note that the operating principle of a ribbon microphone, or the so-called ribbon velocity microphone is by sensing the pressure component, which is fundamentally different than that of a silk or hair-based acoustic flow sensor. The primary sensing element of a ribbon microphone consists of a thin, compliant ribbon which is driven to move from its equilibrium position by the difference in pressure between the two sides under the influence of a sound wave the ribbon (7). The time-varying force on the ribbon of a ribbon microphone will depend on how the acoustic pressure on the two sides of the ribbon differ from each other, which is strongly frequency and geometry dependent as the frequency of the sound increases. The ribbon microphone is unaffected by the viscosity of the fluid while the silk won't experience appreciable acoustic forces in an inviscid medium.

**Highly responsive and tunable acoustic properties of silk threads.** While the macroscale orb-web is more substantial and complex than individual strands of silk, it preserves the unique acoustic responsivity of the individual silk strands. Since the basic element of the orb-web can be regarded as silk strands subjected under boundary conditions, its extraordinary properties to capture the particle velocity of airflow can be partially seen in the acoustic responsivity of an individual strand of silk.

We measured the acoustic responses of a single strand of spider silk under various tensile stress (*SI Appendix*, Fig. S6). The spider silk was supported at both ends. The long axis of the silk (x direction) is perpendicular to the direction of propagation of a harmonic plane wave (z direction). To precisely control the tensile stress within the silk, we bonded one end of spider silk with super glue to one side of a U-shape holder (the plane of U-shape structure is aligned horizontally to the ground) and supported another side of spider silk through the other side of the U-shape holder, with a hanging mass attached at its end. The effective length L of the spider silk was defined by the gap distance of the U-shape holder. The tension T was calculated from the measured weight W of a hanging mass (T=W).

We also created an analytical model to predict the acoustic responsivity of the spider silk, under various tensions. As the sound-induced displacement of the silk is much smaller than the sound wavelength λ in the measured frequency range, the influence of the transverse motion on the variation of the sound field in the z-direction can be ignored. The sound pressure of a harmonic plane sound wave at the frequency ω around the spider silk can be described as $p(t)=Pe^{i\omega t}$, where P is the complex amplitude of the sound pressure, ω=2πf, f is the frequency in Hz. The air particle velocity around the silk can be written as $u(t)= Ue^{i\omega t}$, where $U=V_{air}=P/(\rho_0 c)$ is the complex amplitude of the air particle velocity. The silk is modeled as a string which is subjected to fluid forces by the surrounding medium. The simple approximate analytical model accounting for the transverse motion of a silk can be expressed as,

$$-T\frac{\partial w^2(x,t)}{\partial x^2}+\rho A\frac{\partial w^2(x,t)}{\partial t^2}=f(t) \qquad (1)$$

where w(x, t) is the silk transverse displacement, which depends on both position, x, and time, t, T is the tension of the silk, ρ is the volume density of the material, $A = \pi d^2/4$ is the cross sectional area. The right term f(t) is the aerodynamic force. Considering the problem of a straight cylinder that is moving with some velocity $v(t)= Ve^{i\omega t}$ within a viscous fluid $u(t)= Ue^{i\omega t}$, the forces on this moving cylinder along with the flow field near the cylinder may be written as (8),

$$f(t)=\frac{\rho_0 ckr\pi i}{m}[4\frac{K_1(mr)}{K_0(mr)}+mr]v_r(t) \qquad (2)$$

where $K_0(mr)$ and $K_1(mr)$ are the modified Bessel functions of the second kind of order 0 and 1, respectively, $m=\sqrt{i\omega\rho_0/\mu}$, μ is the dynamic viscosity, $\rho_0$ is the density of the surrounding medium, r = d/2, is the radius of the fiber, $v_r(t)= u(t) - v(t)$ is the relative velocity between the fluid and the cylinder. We may define Z(ω) to be the impedance of the fiber,

$$Z(\omega)=\frac{f(t)}{v_r(t)}=\frac{\rho_0 ckr\pi i}{m}\left[4\frac{K_1(mr)}{K_0(mr)}+mr\right] \qquad (3)$$

To obtain the simplest possible model that accounts for finite boundaries, we assume that the silk of length L is simply-fixed on its ends so that w(0, t)=w(L, t)=0. Equation (1) can be solved as

$$w(x,t)=\sum_{j=1}^{\infty}\frac{Z(\omega)\frac{2}{L}\int_0^L\varphi_j(x)dx}{N\beta_j^2+i\omega Z(\omega)-\omega^2\rho A}u(t)\varphi_j(x) \qquad (4)$$

where j =1, 2, 3, …, ∞ denotes the modal number, $\varphi_j(x)=\sin(n\pi x/L)$ is the modal shape. The fiber velocity $v(x, t) = V(x)e^{i\omega t} = i\omega w(x, t)$, the amplitude ratio of the fiber velocity to the medium velocity $u(t)= Ue^{i\omega t}$ may be expressed as

$$\frac{V(x)}{U}=i\omega\sum_{j=1}^{\infty}\frac{Z(\omega)\frac{2}{L}\int_0^L\varphi_j(x)dx}{N\beta_j^2+i\omega Z(\omega)-\omega^2\rho A}\varphi_j(x) \qquad (5)$$

The analytical model agrees well with the measured results (*SI Appendix*, Fig. S6). Our simplified analytical model and experimental results show that the airborne acoustic responsivity of a single strand of silk could be readily tuned across the tenfold frequency range from hundreds of Hz to thousands of Hz just by applying an equivalent tension produced by a fraction of the weight of the spider body, due to the large length-to-thickness ratio of the spider silk (9). Considering the ultrahigh mechanical strength of the spider silk, the acoustic response of the web threads could be alternated in a wide range of frequencies in real time so as to easily cover various biorelevant frequencies. By adjusting the tension of the web threads, the level and tuning of the mechanical input from the web threads to the metatarsal organ can be both alternated. Meanwhile, as a variably-tensioned network, the orb-web could efficiently filter out the bio-irrelevant low frequency noises which are unavoidable in the natural environment, such as the wind perturbation which has tremendous velocity and pressure amplitude than that of biorelevant acoustic signals.

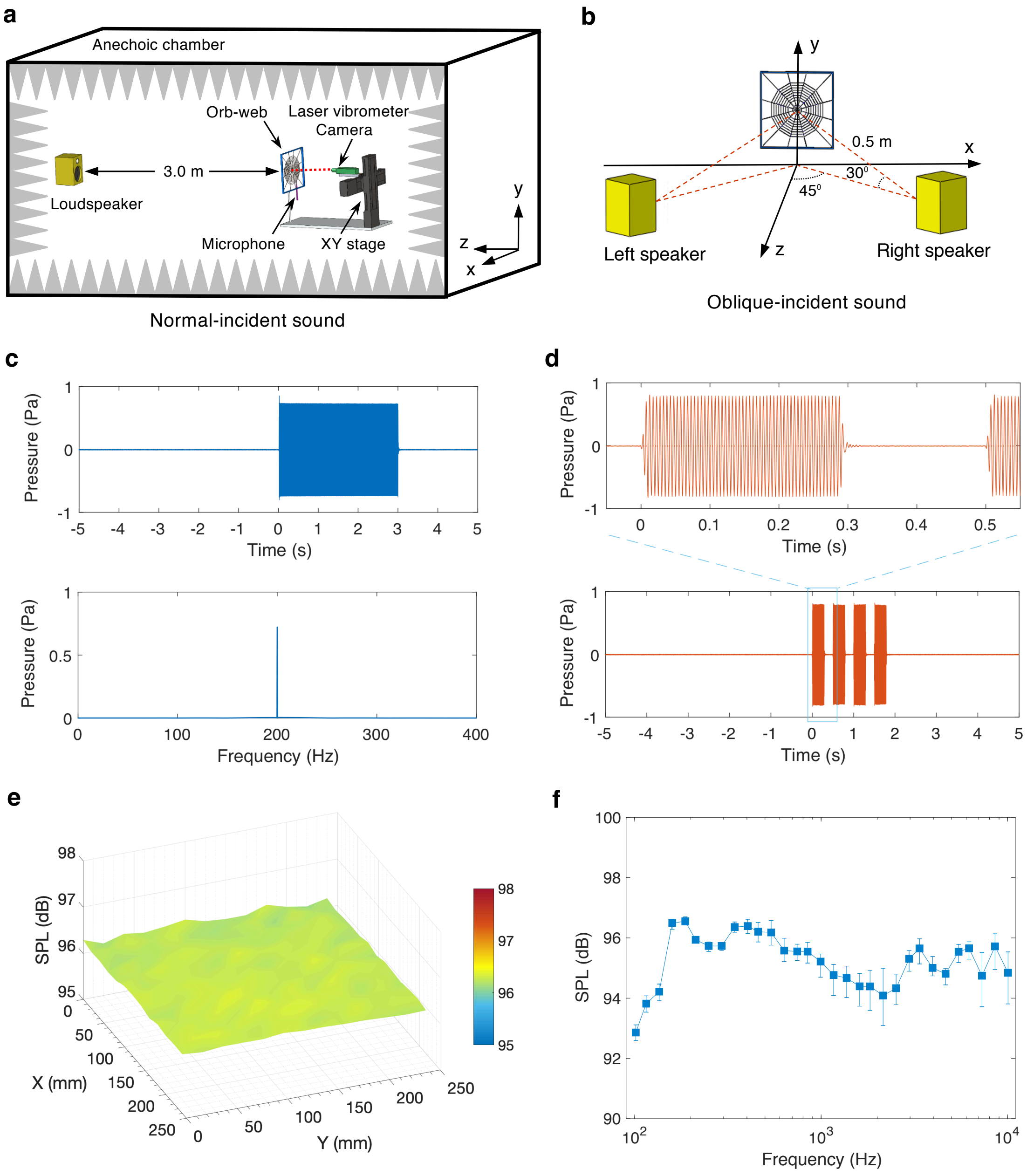


**Fig. S1. Generation of remote airborne sound. a**, **b**, Schematical diagrams of the normal-incident and oblique-incident sound setups. **c**, An example of the normal-incident acoustic stimuli (200 Hz, 88 dB, 3 s duration). The top and bottom images show the time traces and the fast Fourier transform (FFT) of the acoustic signals. **d**, An example of the oblique-incident stimuli. A complete stimulus contains 4 sub directional clips of duration 0.3 s (200 Hz, 88 dB). Individual spider was randomly subjected to one of the patterns of directional acoustic tones, either L+R+L+R or R+L+R+L, where L or R represents the sound generated by the left or right loudspeaker, the symbol (+) represents the silent gap. **e**, An example of the normal-incident sound fields around the spider orb-web at 200 Hz. **f**, Normal-incident sound pressure level (SPL) around the orb-web at a wide range of frequencies. The minimum, mean, and maximum SPL are presented. Normal-incident sound field can be regarded as a planewave approximately.

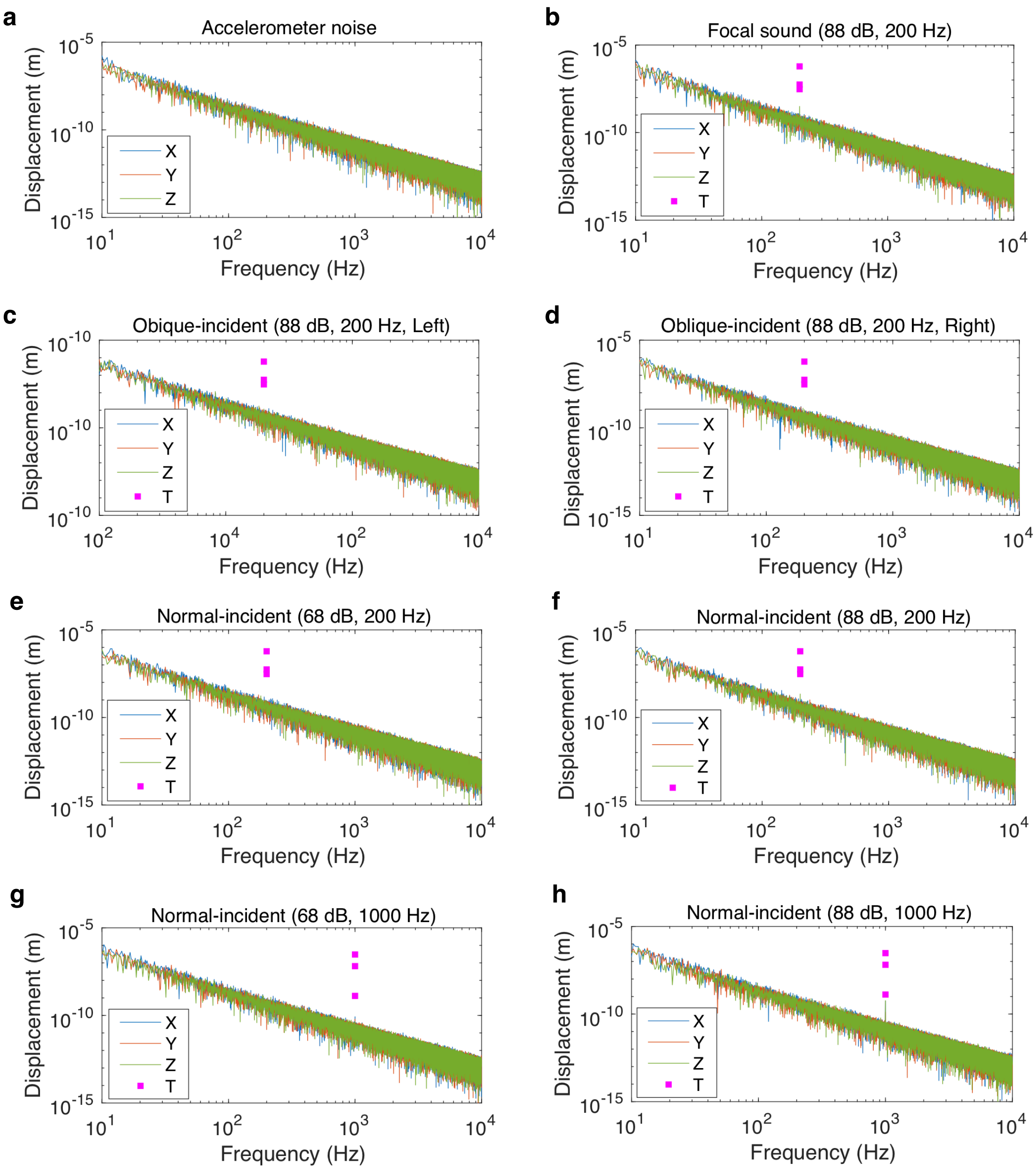


**Fig. S2. Speaker-induced base excitations.** Excitations are measured at the base of the orb-web supporting frame with a tri-axial accelerometer. **a**, Noise floor of the accelerometer. **b**, Base excitations of the focal sound setup. **c**, **d**, Base excitation of the oblique-incident sound setup. **e-h,** Base excitations of the normal-incident sound setup. In the figure legends, X, Y, Z represent the measured 3-dimensional directions, T represents the reported threshold of spider metatarsal lyriform organ (10). Except the excitation condition as shown in **h**, which is more than 2 times (6 dB) lower, all speaker-induced base excitations are more than 10 times (20 dB) lower than the threshold of the spider metatarsal lyriform organ.

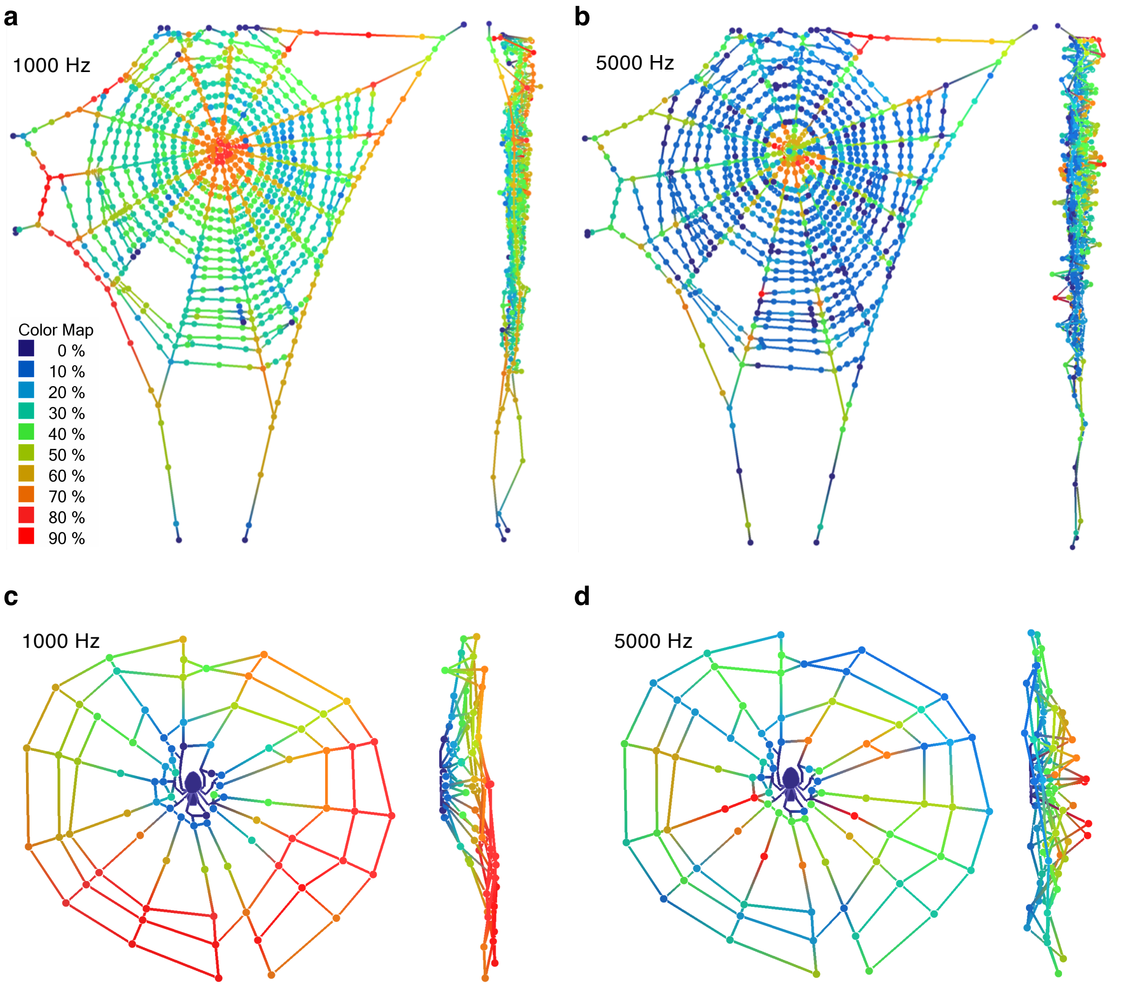


**Fig. S3. Orb-web responses to remote normal-incident sound. a**, **b**, Out-of-plane motion of a complete orb-web induced by a steady-state sound field at 1000 Hz and 5000 Hz. **c, d**, Out-of-plane motion of an orb-web containing a live spider, induced by a sound field at 1000 Hz and 5000 Hz. Color coding of **b, c, d** same as **a**. The colored heat map represents the amplitude ratio of the web thread velocity V to the air particle velocity $V_{air}$.

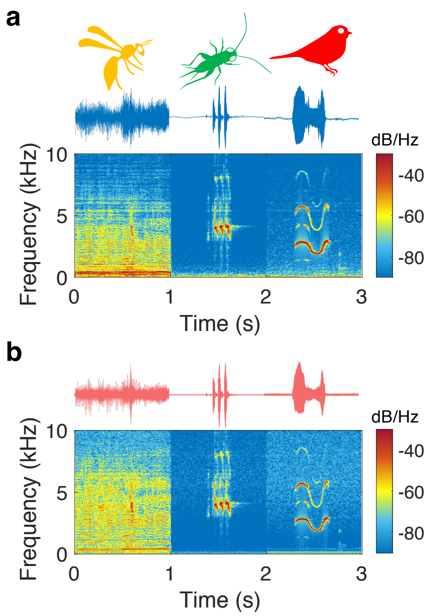


**Fig. S4. Time-domain traces and spectrograms of the sound-induced orb-web motion containing a live spider.** The normal-incident airborne acoustic signal (**a**) was measured by a pressure microphone nearby the spider. The out-of-plane web motion (**b**) was measured by a laser vibrometer at a radial web thread, 2 mm away from the spider foreleg.

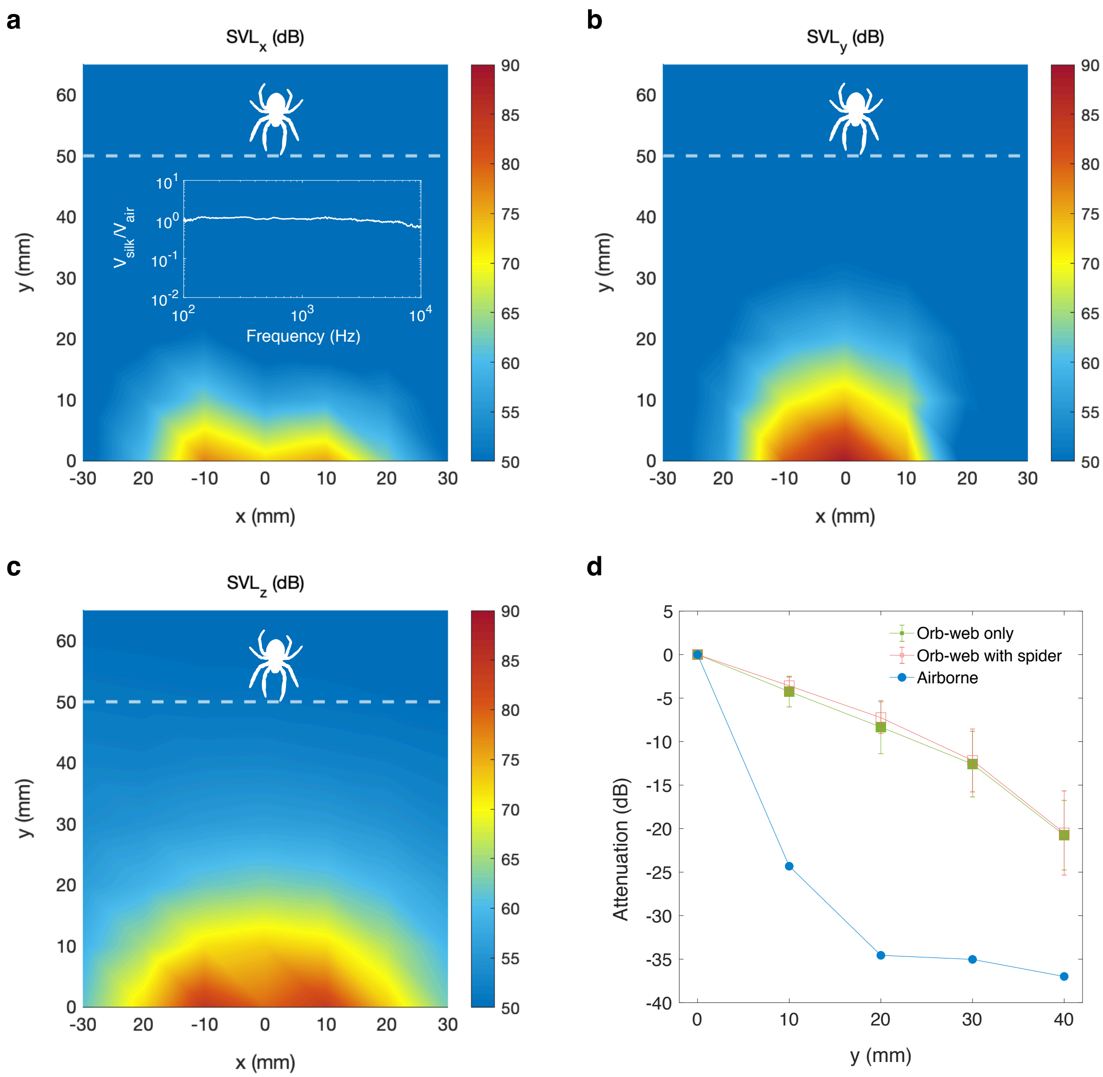


**Fig. S5. Focal airborne acoustic field generated by the miniature speaker. a-c,** Measured sound velocity level (SVL) in 3-dimensions along the plane of the orb-web. The air particle velocity in each dimension was mapped by a silk velocity probe, whose characteristics are shown as the insert figure in **a**. The 3-dimensional velocity components constitute the overall SVL as shown in Fig. 3*C*. **d**, Statistic attenuation of the out-of-plane web vibration along the radial threads of orb-webs (N=12 spiders). Error bars show one standard deviation. The near-field airborne signal is also shown as comparison, extracted from the overall SVL measured at multiple locations (Fig. 3*C*, x=0 mm, y=0~40 mm). The out-of-plane motion of web threads induced by the mini-speaker attenuated slower than the airborne signals, so as to transmit the vibrational signals to the spider.

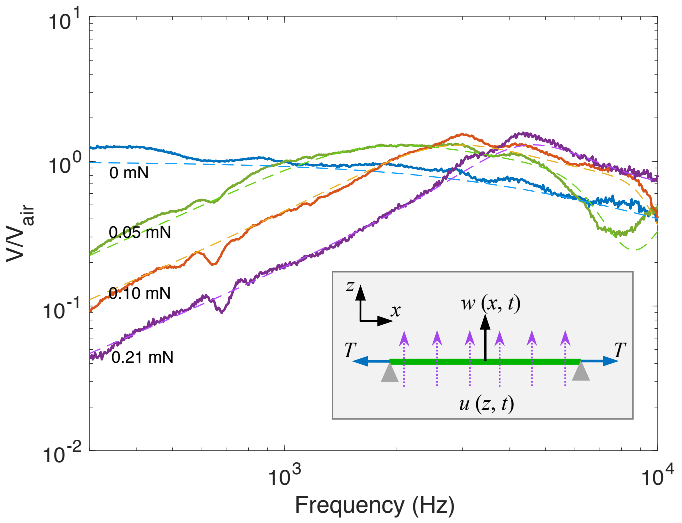


**Fig. S6. Tuning the acoustic properties of the spider silk by tension.** The solid lines show the frequency responses of a single strand of spider silk (L=3 cm, d=1.5 μm) measured in the middle position under various tensile stress. The spider silk is supported at its two ends as shown by the insert figure. The propagation direction of the acoustic waves is perpendicular to the direction of the silk. The dash lines show the predicted frequency responses according to our simplified analytical model (see Supplementary Text). The applied tension (0-0.21 mN) ranges from zero to a fraction of the body weight of spiders (0.7-1.9 mN).

**Table S1. Experimental configurations for spider behavioral testing.**

| Sound type | Speaker type | Distance (m) | Frequency (Hz) | SPL/SVL (dB) | Duration (s) | Spider Number |
|---|---|---|---|---|---|---|
| Normal-incident sound | Loudspeaker | 3.0 | 200 | 68 | 3 | 12 |
| | | | 200 | 88 | 3 | 12 |
| | | | 1000 | 68 | 3 | 12 |
| | | | 1000 | 88 | 3 | 12 |
| Oblique-incident sound (L/R) | Loudspeaker | 0.5 | 200 | 88 | 0.3x4 | 12 |
| Focal sound | Mini-speaker | 0.050 | 200 | ≤ 68 | 3 | 12 |

**Table S2. Sound loudness produced by distant animals.** For the comparison of sound loudness between different species measured at various distances, the sound pressure level (SPL) of remote songs measured at a distance (d) is converted to SPL at 10 m distance according to the inverse square law for acoustics, $SPL_{10m} = -20\log_{10}(10/d) + SPL_{d}$. Meanwhile, the air particle velocity ($V_{air}$) of local wing beats at a measured position is converted to sound velocity level (SVL). With a hearing threshold lower than 68 dB, orb-weaving spiders (*Larinioides sclopetarius*) should be able to perceive the remote sounds produced by various animals such as birds and crickets at a distance more than 10 m away.

| **Sound type** | **Animal Species** | **Distance (m)** | **SPL/SVL (dB)** |
|---|---|---|---|
| Remote song | White bellbird (*Procnias albus*) (11) | 10 | 105 |
| | Field cricket (*Gryllus campestris*) (12) | 10 | 80 |
| | Shrill Thorntree Cicada (*Brevisana brevis*) (13) | 10 | 81 |
| | Emerald forest frog (*Hylorina sylvatica*) (14) | 10 | 83 |
| Local wing beat | Median wasp (*Dolichovespula media*) (15) | 0.1 | >100 dB |
| | Blowfly (C*alliphora erythrocephala*) (16) | 0.1 | 90 |
| | Stingless bees (*Melipona scutellaris*) (17) | 0.02 | 89 |
| | Southern house mosquito (*Culex quinquefasciatus*) (18) | 0.03 | 72 |

**Table S3. Spider behavioral responses to successive presentations of the oblique-incident sound stimulus.** The stimuli (200 Hz, 88 dB) are either L+R+L+R or R+L+R+L, where L and R represents the stimulus from the left or right direction with 0.3 s duration, the symbols (+) represent silent gaps, which are 0.2 s, 1 s, and 0.2 s respectively. Behavioral responses 1) before the first stimulus, 2) during each sub stimulus, and 3) after the last stimulus are listed. In the table, blank represents no response, *Sit* represents that a spider resting back to the initial hub region after stimulus, *Fall* represents that a spider falling off the orb-web.

| Spider No. | Before stimulus | Sub-1 stimulus | Sub-2 stimulus | Sub-3 stimulus | Sub-4 stimulus | After stimulus |
|---|---|---|---|---|---|---|
| 1 | | *Turn* | *Raise leg* | *Raise leg* | *Raise leg* | *Sit* |
| 2 | | | | | *Turn* | *Sit* |
| 3 | | | | | | Crouch |
| 4 | | | | | | *Crouch* |
| 5 | | *Stretch* | | | | *Sit* |
| 6 | | | *Raise leg* | *Raise leg* | *Raise leg* | *Sit* |
| 7 | | *Turn* | *Turn* | *Turn* | *Turn* | *Sit* |
| 8 | | | | | | *Raise leg, Sit* |
| 9 | | *Turn* | *Raise leg* | *Raise leg* | *Raise leg* | *Sit* |
| 10 | | | | *Stretch* | *Stretch* | *Sit* |
| 11 | | *Turn* | *Turn, Fall* | | | |
| 12 | | | | | | *Crouch* |

**Movie S1 (separate file). Spider behavioral responses to distant (3 m) airborne sound.** Example behaviors are recorded under 200 Hz normal-incident airborne acoustic tones at 88 dB. In the first part, spider lifts its forelegs into air. After the initial response, spider shakes its orb-web. In the second part, spider turns its body instantaneously. Spider forelegs are afterwards stretched into air. In the third part, spider extends legs out of its body direction instantaneously. As an incidental action, spider forelegs are stretched into air. In the last part, spider pulls the web threads slightly towards its body direction with legs.

**Movie S2 (separate file). Spider source localization.** Example of spider behavioral responses to oblique-incident (L/R, 45° in azimuth, 0.5 m to the spider) acoustic waves is presented. Spider turns its body towards the sound sources in response to the first and second sub-audio clips. Afterwards, spider lifts its forelegs in response to the third and last sub-audio clips.

**Movie S3 (separate file). Out-of-plane motion of the spider and its orb-web induced by normal-incident (3 m) airborne sound.** In the first part, motion of a complete orb-web without the spider resting in the hub is shown. In the second part, motion of a spider and its peripheral orb-web is shown. The colored heat map represents the amplitude ratio of the silk thread (or spider body) velocity V to the air particle velocity $V_{air}$. The plane of the orb-web is tilted by 20° for visualization of the out-of-plane motion.

**Audio S1 (separate file). Measured orb-web motion induced by distant (3 m) normal-incident airborne sound.** Orb-web motion signals were measured by laser Doppler vibrometer at the hub region. The airborne signals contain broadband acoustic information, including wing beats of insects, cricket calls and bird songs.

### SI References


1. P. S. Shamble, et al., Airborne acoustic perception by a jumping spider. *Curr. Biol.* **26**, 2913–2920 (2016).
2. G. Menda, et al., The long and short of hearing in the mosquito *Aedes aegypti*. *Curr. Biol.* **29**, 709-714.e4 (2019).
3. M. Gregorič, I. Agnarsson, T. A. Blackledge, M. Kuntner, Darwin's bark spider: giant prey in giant orb webs (*Caerostris darwini*, Araneae: Araneidae)? *J. Arachnol.* **39**, 287–295 (2011).
4. C. T. F. Ross, T. late J. Case, A. Chilver, Strength of Materials and Structures (Elsevier, 1999).
5. S. F. Hoerner, Fluid-dynamic Drag (Hoerner Fluid Dyn, 1965).
6. F. G. Omenetto, D. L. Kaplan, New opportunities for an ancient material. *Science* **329**, 528–531 (2010).
7. H. F. Olson, Acoustical Engineering (Van Nostrand, 1957).
8. R. N. Miles, J. Zhou, Sound-induced motion of a nanoscale fiber. *J. Vib. Acoust.* **140** (2017).
9. J. Zhou, N. Moldovan, L. Stan, H. Cai, D. A. Czaplewski, and D. López, Approaching the strain-free limit in ultrathin nanomechanical resonators. *Nano Lett*., 20, 5693-5698 (2020).
10. F. G. Barth, Geethabali, Spider vibration receptors: Threshold curves of individual slits in the metatarsal lyriform organ. *J. Comp. Physiol.* **148**, 175–185 (1982).
11. J. Podos, M. Cohn-Haft, Extremely loud mating songs at close range in white bellbirds. *Curr. Biol.* **29**, R1068–R1069 (2019).
12. H. Nocke, Physiological aspects of sound communication in crickets (*Gryllus campestris L.*). *J. Comp. Physiol.* **80**, 141–162 (1972).
13. Villet Martin, Sound pressure levels of some African cicadas (*Homoptera*: *Cicadoidea*). *J. Entomol. Soc. South. Afr.* **50**, 269–273 (1987).
14. M. Penna, R. Solís, Frog call intensities and sound propagation in the South American temperate forest region. *Behav. Ecol. Sociobiol.* **42**, 371–381 (1998).
15. J. Tautz, H. Markl, Caterpillars detect flying wasps by hairs sensitive to airborne vibration. *Behav. Ecol. Sociobiol.* **4**, 101–110 (1978).
16. C. Klopsch, "The flow field around a flying blowfly: characteristics and guidance of spider prey capture behavior." (Doctoral Dissertation, 2010).
17. M. Hrncir, D. L. P. Schorkopf, V. M. Schmidt, R. Zucchi, F. G. Barth, The sound field generated by tethered stingless bees (*Melipona scutellaris*): inferences on its potential as a recruitment mechanism inside the hive. *J. Exp. Biol.* **211**, 686–698 (2008).
18. T. Nakata, et al., Aerodynamic imaging by mosquitoes inspires a surface detector for autonomous flying vehicles. *Science* **368**, 634–637 (2020)
